# The sound of restored soil: Measuring soil biodiversity in a forest restoration chronosequence with ecoacoustics

**DOI:** 10.1101/2023.01.23.525240

**Authors:** Jake M. Robinson, Martin F. Breed, Carlos Abrahams

## Abstract

Forest restoration requires monitoring to assess changes in above- and below-ground communities, which is challenging due to practical and resource limitations. With emerging sound recording technologies, ecological acoustic survey methods—also known as ‘ecoacoustics’—are increasingly available. These provide a rapid, effective, and non-intrusive means of monitoring biodiversity. Above-ground ecoacoustics is increasingly widespread, but soil ecoacoustics has yet to be utilised in restoration despite its demonstrable effectiveness at detecting meso- and macrofauna acoustic signals. This study applied ecoacoustic tools and indices (Acoustic Complexity Index, Normalised Difference Soundscape Index, and Bioacoustic Index) to measure above- and below-ground biodiversity in a forest restoration chronosequence. We hypothesised that higher acoustic complexity, diversity and high-frequency to low-frequency ratio would be detected in restored forest plots. We collected *n* = 198 below-ground samples and *n* = 180 ambient and controlled samples from three recently degraded (within 10 years) and three restored (30-51 years ago) deciduous forest plots across three monthly visits. We used passive acoustic monitoring to record above-ground biological sounds and a below-ground sampling device and sound-attenuation chamber to record soil communities. We found that restored plot acoustic complexity and diversity were higher in the sound-attenuation chamber soil but not *in situ* or above-ground samples. Moreover, we found that restored plots had a significantly greater high-frequency to low-frequency ratio for soil, but no such association for above-ground samples. Our results suggest that ecoacoustics has the potential to monitor below-ground biodiversity, adding to the restoration ecologist’s toolkit and supporting global ecosystem recovery.

**Implications for Practice:**

- This is the first known study to assess the sounds of soil biodiversity in a forest restoration context, paving the way for more comprehensive studies and practical applications to support global ecosystem recovery.
- Soil ecoacoustics has the potential to support restoration ecology/biodiversity assessments, providing a minimally intrusive, cost-effective and rapid surveying tool. The methods are also relatively simple to learn and apply.
- Ecoacoustics can contribute toward overcoming the profound challenge of quantifying the effectiveness (i.e., success) of forest restoration interventions in reinstating target species, functions and so-called ‘services’ and reducing disturbance.

## Introduction

In the absence of large-scale ecosystem restoration and effective monitoring strategies, 95% of the Earth’s land is projected to be degraded by 2050 (Yu et al. 2020). This includes forests—ecosystems that comprise a combination of species, geology and climatic processes in which trees are the dominant vegetation type (Kimmins 2004; Glatthorn et al. 2021; Seidl and Turner 2022). The integrity of forest ecosystems depends on a rich tapestry of biodiversity (Müller 2000; Watson et al. 2018). Microscopic organisms or ‘microbiota’ provide forest trees with nutrients and the ability to communicate via mycorrhizae (Simard 2018; Robinson et al. 2021), and soil meso- and macrofauna contribute to soil formation and energy flows (Le Bayon et al. 2021). The strength and complexity of the relationships between organisms confer resilience to forest ecosystems. Without this complexity, the integrity of forests diminishes, and their capacity to respond to environmental stressors, such as extreme heat caused by climate change, is inhibited (Messier et al. 2019; Pardos et al. 2021). Deforestation—the purposeful clearing of forested land—now occurs at a rapid pace globally. Indeed, the tropics alone lost 12.2 million ha of tree coverage in 2020, an area three times the size of the Netherlands (Sama 2021; Gola et al. 2022). Deforestation contributes to global species extinctions, which are currently occurring at 1,000 times higher than the natural background rate (De Vos 2015). Deforestation also reduces key functional elements (so-called ‘ecosystem services’) that benefit humans, such as stormwater management, climate regulation, sustainable resources and recreational amenities (Li et al. 2007; Taye et al. 2021). Therefore, effective forest restoration strategies are vital to biodiversity and human wellbeing.

Forest restoration is often conceptualised as intervening to convert a degraded forest starting point to an endpoint that is an idealised natural forest, whilst recognising that restoring functions is a priority (Stantfurt et al. 2014). However, a profound challenge in this process is quantifying the effectiveness (i.e. success) of forest restoration interventions in reinstating target species, functions and ‘services’ (Camarretta et al. 2020), and reducing further disturbance. Indeed, ecosystem restoration can be viewed as a continuum of stages from planning to implementation to monitoring (Robinson et al. 2022). The monitoring stage plays a crucial role in quantifying the effectiveness of restoration interventions by measuring recovery and potential ongoing disturbance (de Almeida et al. 2020). Primary observations and derived measurements of changes in biodiversity status are considered fundamental to monitoring the effectiveness of restoration strategies (Breed et al. 2019; Hansen et al. 2021). This is exemplified by GEO BON Essential Biodiversity Variables (EBVs), which provide the first level of abstraction between low-level observations and high-level indicators of biodiversity (Kissling et al. 2018). However, acquiring these EBVs, which include genetic composition, species populations, species traits, community composition, ecosystem functioning and ecosystem structure (O’Connor et al. 2020), via traditional survey methods can be time and resource-intensive and potentially intrusive (Gollan et al. 2013; Beng et al. 2020; Hoban et al. 2022).

Due to these constraints, forest restoration data are often limited to visible macro-organisms, particularly the trees and other floral and faunal assemblages above-ground (Stoddard et al. 2011; Williams-Linera et al. 2021). Moreover, ecological data are often ambiguous and, therefore, incompatible with further research (Zipkin et al. 2021). With the advent of new sound recording technologies, ecological acoustic survey methods, also known as ‘ecoacoustics’, are becoming increasingly available (Abrahams and Geary 2020; Abrahams et al. 2021; Müller et al. 2022). They can provide effective and non-invasive approaches to gathering biodiversity data—e.g. on target species, assemblages and environmental variables essential to restoration monitoring (Teixeira et al. 2019; Stowell and Sueur 2020). In recent years, ecoacoustics has been applied to monitor elusive species in several environmental contexts—particularly in conservation biology (Teixeira et al. 2019; Stowell and Sueur 2020). For instance, passive acoustic monitoring (often shortened to ‘PAM’), which involves deploying autonomous acoustic sensors, has been used to collect recordings of biological sounds (known as ‘biophony’) from bats (Hintze et al. 2021; López-Baucells et al. 2021), birds (Abrahams 2019; Abrahams and Geary 2021), and invertebrates (Harvey et al. 2011; van der Mescht et al. 2021; Mankin et al. 2022) in terrestrial environments; and cetaceans (Jones et al. 2020; Guidi et al. 2021), amphibians (Gan et al. 2020), crustaceans (Kühn et al. 2022), and fish (Popper and Hawkins 2019) in aquatic environments. Indeed, ecoacoustics has emerged as an efficient tool to measure and monitor biodiversity and has the potential to enhance the toolbox of restoration ecologists. Moreover, the same audio recording devices can detect anthropogenic noise (known as ‘anthrophony’) (de Framond and Brumm 2022). Anthrophony may contribute to ecosystem degradation by adversely affecting animal fitness, health (De Jong et al. 2018; Kleist et al. 2018) and behaviour (Tidau and Briffa 2019; Hastie et al. 2021), and the composition and functionality of microbial communities (Robinson et al. 2021). Therefore, ecoacoustics could provide important measurements across the degradation-restoration continuum.

Despite the potential of ecoacoustics to contribute to forest restoration monitoring, few studies have deployed this technology to assess above-ground faunal soundscapes in a forest restoration context (Turner et al. 2018; Vega-Hidalgo et al. 2021). Moreover, to our knowledge, no studies have applied ecoacoustics to measure or monitor below-ground biodiversity in a restoration context. This is despite its demonstrable effectiveness at detecting soil meso- and macrofauna acoustic signals in other settings, such as agriculture (Maeder et al. 2019), silviculture (Maeder et al. 2022), and in controlled chambers (Lacoste et al. 2018). Here we apply novel ecoacoustics devices to measure above- and below-ground biodiversity in a forest restoration chronosequence (a set of ecological sites that share similar attributes but represent different times since restoration), using a range of acoustic indices to analyse the recordings, including the Acoustic Complexity Index (ACI) (Pieretti et al. 2011), Normalised Difference Soundscape Index (NDSI) (Kasten et al. 2012), and Bioacoustic Index (BI) (Boelman et al. 2007). As faunal species richness, abundance, biomass and functional diversity are known to increase with restoration age (Derhé et al. 2016), we expected acoustic diversity to increase accordingly. Specifically, our study aimed to test the following hypotheses:

a. Acoustic complexity/diversity will be higher in restored plots (30-50 years since restoration), compared with degraded plots (0-10 years since clearing without any active restoration intervention), in both soil and ambient recordings.
b. The high-frequency to low-frequency ratio (an amended version of the Bioacoustic Index) will be higher in restored plots than in degraded plots. This would indicate lower noise disturbance in the restored plots, based on the assumption that high-frequency sounds are more representative of biophony than low-frequency anthrophony resulting from mechanical noise and ground vibrations.
c. Soil acoustic diversity will positively correlate with invertebrate abundance and richness, with higher scores in the restored plots.

## Materials and Methods

### Study location

Greno Woods is a large forest (169 ha) near Sheffield in South Yorkshire, UK (Fig. 1). The forest comprises several restoration age classes. Due to comparator site availability constraints, samples were collected from two age classes: 0-10 years since deforestation and no active restoration interventions since (referred to in this study as ‘degraded’) representing recent degradation; and 31-50 years since restoration (referred to in this study as ‘restored’). We identified three spatially-independent replicate plots for sampling each age class (0A, 0B, 0C, and 30A, 30B, 30C; Fig. 1 and Fig. 2A, B). The habitat classification of all restored sampling plots was semi-natural broadleaved woodland of the W16 National Vegetation Classification (Rodwell 2006). The degraded plots were dominated by bracken *Pteridium aquilinum*, with occasional silver birch *Betula pendula* saplings. The restored plots were dominated by English oak *Quercus robur*, sessile oak *Q. petraea*, silver birch and rowan *Sorbus aucuparia*, with a well-developed understory of bilberry *Vaccinium myrtillus*, bramble *Rubus fruticosus agg*., holly *Ilex aquifolium*, and bracken.

**Figure 1.**
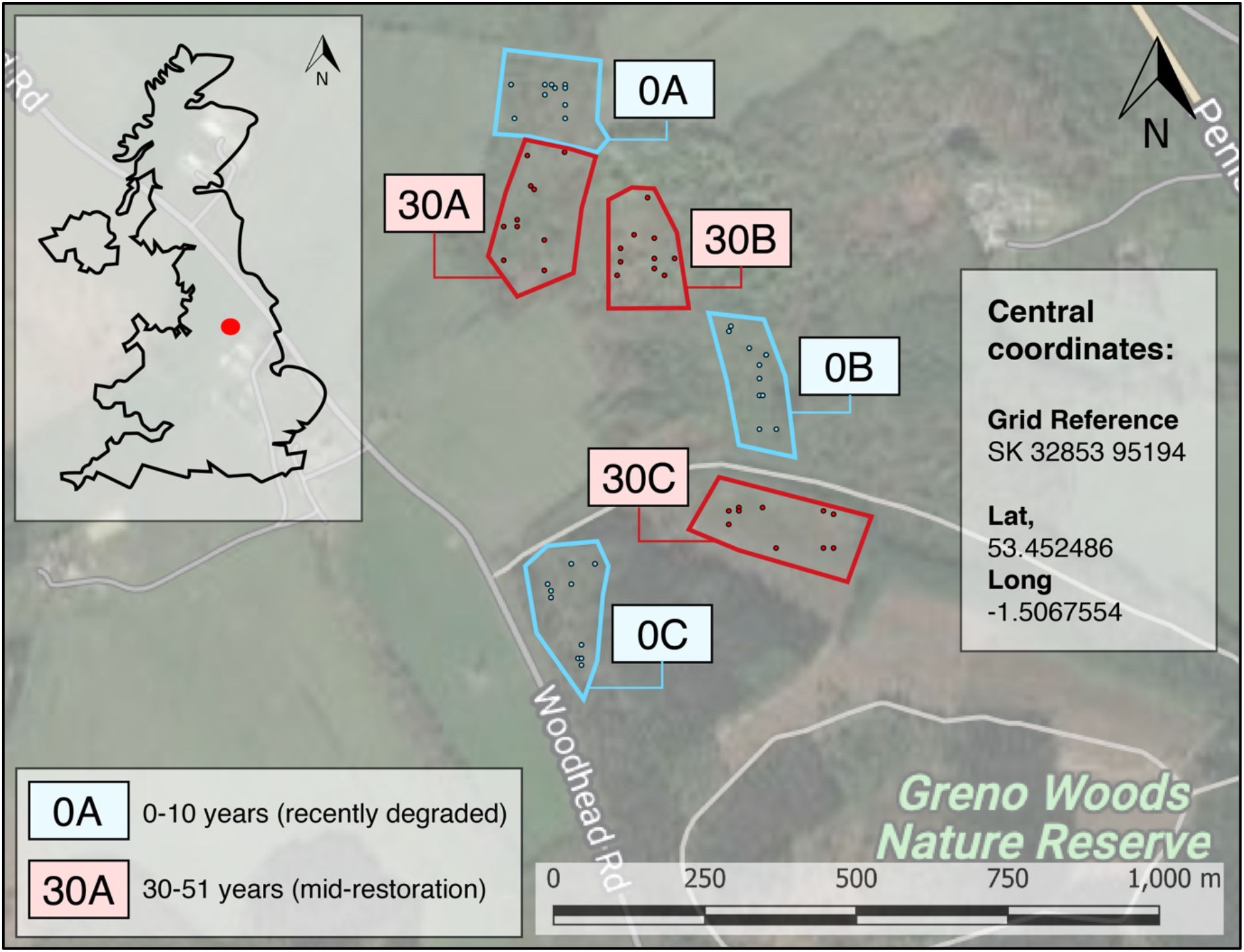
Site location (Greno Woods, South Yorkshire, UK), sampling plots within the blue polygons for degraded and red polygons for restored, and the ten randomly selected sampling locations within each plot. The inset shows the study location (red dot) in the broader UK context.

**Figure 2.**
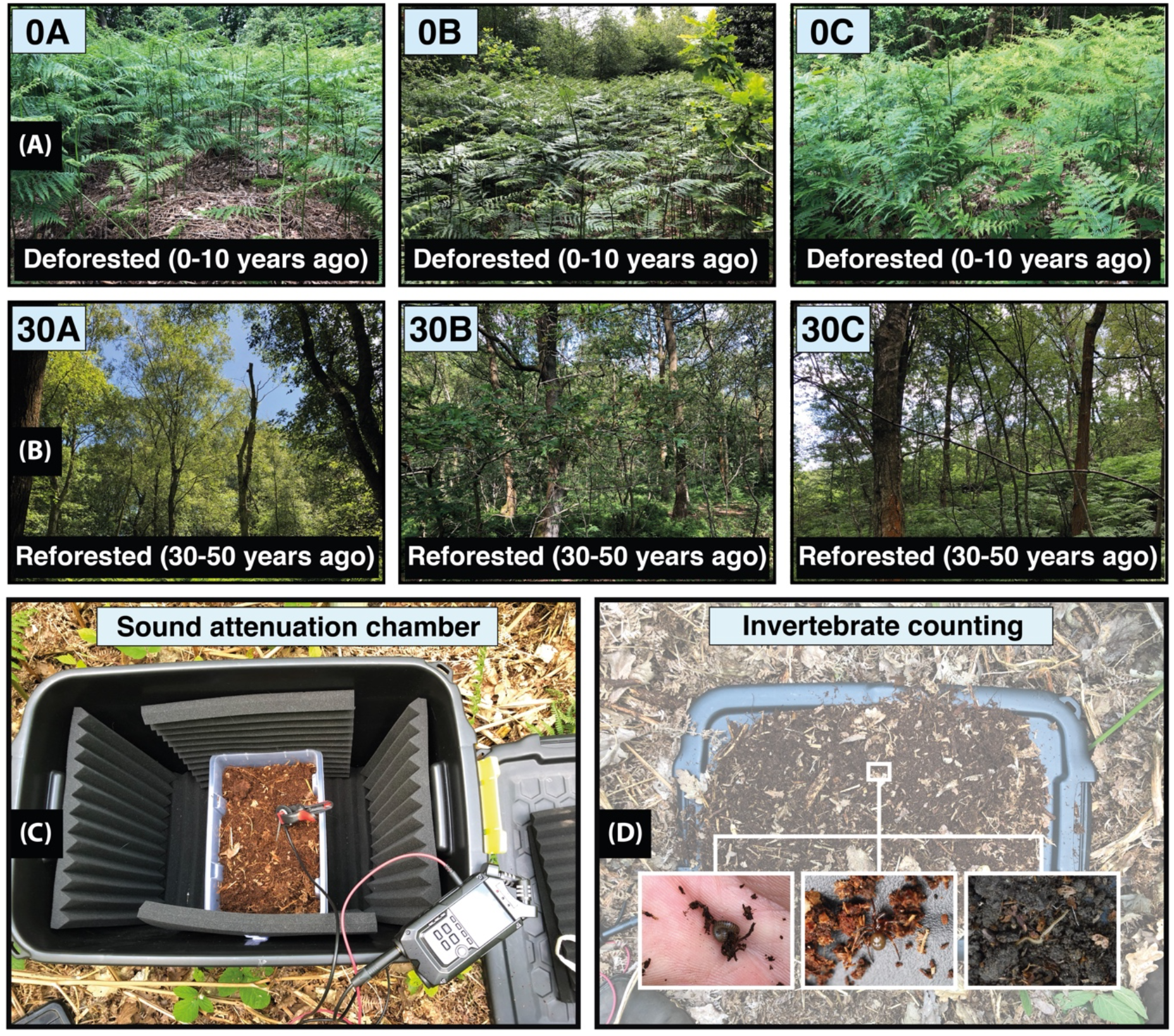
(A) Degraded study plots. (B) Restored study plots. (C) Sound attenuation chamber with the Zoom H4n recorder and JrF C-series contact microphone. (D) The invertebrate counting method.

### Soundscape sampling

We used a relatively inexpensive ecoacoustics sampling device for below-ground sampling: a JrF C-Series Pro contact microphone sensor (jezrileyfrench.co.uk) with a 2 m cable and a 1/4” Neutrik jack. The C-series contact microphones provide a broader frequency response than others, meaning more low-end and mid-frequency range responses. This broader frequency response is optimal for recording below-ground soundscapes (Maeder et al. 2019; Gamal et al. 2020). The JrF microphone was attached to a metal probe and linked to a handheld acoustic recording device (Zoom H4n Pro) prior to inserting the probe into the soil. We recorded .wav sound files, at 16 bit depth, and with a sampling rate of 48 kHz, which is a similar rate used in other soil acoustic research (Abrahams 2019), capturing sounds to a maximum of 24 kHz and therefore covering the entire audible range (Maeder et al. 2022). To record above-ground (ambient) sound—for instance, to detect soniferous species such as birds—we installed a Tascam DR-100MKII audio recording device onto a tripod in each plot, using its inbuilt omni-directional microphones to record sounds with the same file format.

We selected below-ground acoustic sampling locations using a geographical information system (GIS). We created polygon boundary shapefiles around each of the six spatially-independent sampling plots and generated ten random sampling points for each plot using the random points algorithm in QGIS (version 3.24.3 ‘Tisler’). Below-ground sound samples were collected from the predetermined random points within each plot. We repeated the sampling on three occasions across three months (June, July, and August) in the summer of 2022 (Table S1).

To determine the appropriate sampling duration for below-ground samples, we first ran a pilot study, testing the potential saturation and decay of acoustic indices using different sampling durations (20 s, 1 min, 3 mins, and 5 mins). The sampling durations were randomised and collected over two visits (*n* = 14 per sampling duration). Each recording followed a separate probe insertion into the soil to represent the main study approach. To control for initial geophony (e.g., displaced soil particles) and potential disturbance to biophony from the physical disturbance of entering the soil, recordings always followed an initial 30 s resting period. We also controlled for higher frequency anthrophony by setting a low-pass filter to 2 kHz during analysis. This testing process identified a sampling duration of 3 mins as optimal. There was no significant effect of time post-3 mins (i.e. 3 mins vs. 5 mins) on acoustic complexity (t = 1.7-2.1; *p* = 0.48-0.64) (Fig. S1).

Following the pilot study, we collected data for the main part of the study. During the three sampling occasions, we set the Tascam DR-100MKIII to record above-ground soundscape samples at each plot. We then recorded the 3 mins below-ground samples (*n* = 10) in each plot, alongside simultaneous control samples of the same duration. The latter involved recording ‘blanks’ by leaving a recorder and contact microphone outside the soil, supported on sound attenuation foam. In total, we collected *n* = 180 below-ground samples (3 mins each) with their matching control recordings, and *n* = 18 above-ground samples. The above-ground recordings were post-processed by being divided into 3 mins sections to simultaneously match the below-ground recordings (*n* = 180 subsamples).

### Sound attenuation chamber

We used an additional sampling method to record the soil soundscape in each plot. This involved collecting soil samples with a 3L plastic container and placing them into a sound-attenuation chamber, allowing us to record a ‘snapshot’ of the soundscape under controlled conditions (Fig. 2C). We used the same recording equipment for the *in situ* and sound-attenuation chamber samples. In total, we collected *n* = 18 chamber samples (3 mins each). To determine the optimal sound-attenuation chamber design, we first ran a pilot study using different sound barrier configurations (Fig. S2). The final design comprised a 60 L plastic chamber, with sound-attenuation foam installed on each internal wall, including the base and lid (Fig. 2C).

### Invertebrate counts

We recorded the abundance and richness of meso- and macrofauna in the soil by collecting 3 L soil samples from a random point (determined using a digital number randomiser). We subsequently counted the invertebrates on the sound-attenuation chamber lid (Fig. 2D) by systematically searching through the soil, working from left-to-right and carefully displacing soil particles, thereby revealing the invertebrates (Stroud 2019). The invertebrates were photographed and recorded in a spreadsheet on-site. The soil and the invertebrates were placed back in their source location once the counting was completed.

### Data analysis

To process the sound recordings (.wav files), we used the wildlife sound analysis software Kaleidoscope Pro (Version 5.4.7; Wildlife Acoustics, 2022). This software allows for the analysis of full-spectrum recordings to measure multiple acoustic indices, including the ACI (Pieretti et al. 2011), NDSI (Kasten et al. 2012) and BI (Boelman et al. 2007) selected for this study. We chose two diversity indices (ACI and BI) and one index to measure the biophony-to-anthrophony ratio (NDSI), allowing us to test our three hypotheses.

ACI directly measures the variability in sound intensity in both frequency and time domains, comparing the normalised absolute difference of amplitude between adjacent FFT windows in each frequency bin over a period of K seconds. First, it computes the absolute difference between adjacent values of intensity:

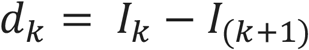

The changes in the recording’s temporal step are encompassed by the summation of the *d*″:

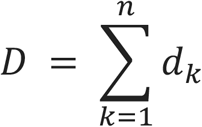

To obtain the relative intensity and reduce the influence of the distance between the microphone and biophony source, the result *D* is divided by the total sum of the intensity values (Maeder et al. 2022):

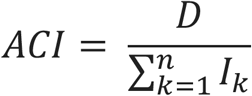

The total ACI is the sum of the ACIs across bins for each period K in the recording.

BI is computed as “the area under each curve including all frequency bands associated with the dB value that was greater than the minimum dB value for each curve. The area values are thus a function of both the sound level and the number of frequency bands” (Boelman et al. 2007).

NDSI is computed as follows:

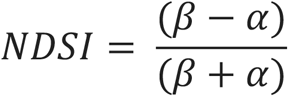

Where *β* and *α* are the total estimated power spectral density for the largest 1 kHz biophony bin and the anthrophony bin, respectively. The NDSI is a ratio in the range [− 1 to + 1], where + 1 indicates a signal containing only high-frequency biophony and no low-frequency anthrophony (Kasten et al. 2012).

Standard settings in Kaleidoscope Pro were used for the calculation of above-ground acoustic indices. However, as sounds above 2 kHz do not propagate well through the soil (Maeder et al. 2022), for the below-ground acoustic indices, we set a maximum frequency of 2 kHz, and a lower threshold of 500 Hz for biophony in NDSI and BI.

Standard settings in Kaleidoscope Pro were used for the calculation of above-ground acoustic indices. However, as sounds above 2 kHz do not propagate well through the soil (Maeder et al. 2022), for the below-ground acoustic indices, we set a maximum frequency of 2 kHz, and a lower threshold of 500 Hz for biophony in NDSI and BI.

All statistical analysis was conducted in R Version 2022.02.2 ‘Prairie Trillium’ (R Core Team 2022) with supplementary software (e.g., Microsoft Excel for .csv file processing). To test for the effect of restoration on acoustic index values, we applied the two-samples t-test using the rstatix package (Kassambara 2022). We also fit linear mixed effects models (LMM) to the data using R and its lme4 package (Bates et al. 2015), with separate models fitted for different plots and visits. LMMs included random effects (plots and visits), which are essential to account for the spatial and temporal correlation between the plots and visits in our experimental design. Acoustic index outputs were included as response variables, and the degraded vs. restored plots were included as fixed effects (predictor variables). Tests of significance were conducted using Satterthwaite’s degrees of freedom t-test, which is a function of the LmerTest package in R (Kuznetsova 2020). Soil invertebrate beta diversity was visualised using nonmetric multidimensional scaling (NMDS) ordination of Bray–Curtis distances using the Vegan package in R (Oksanen et al. 2022). The ordination plots show low-dimensional ordination space in which similar samples are plotted close together, and dissimilar samples are plotted far apart. We used the analysis of similarities (ANOSIM) approach to test for compositional differences between treatment groups. Data visualisations were produced using a combination of R and the Adobe Illustrator creative cloud 2021 version (Adobe 2021).

## Results

### Soil invertebrate observational surveys

#### Restored/degraded soil invertebrate abundance and richness

Restored soils had higher invertebrate abundance (t-test: t = −2.2, df = 8, *p* = 0.02), and there was no significant effect of restoration/degradation status on invertebrate richness (t = 0, df = 8, *p* = 1).

#### Beta diversity

Soil invertebrate community composition was significantly different between degraded and restored plots (stress 0.01, R: 0.55, *p* = 0.05, permutations = 999) (Fig. 6). Earthworms (sub-order: Lumbricina) were the dominant invertebrate in the soil for both treatment groups (*n* = 13 from *n* = 64 for degraded vs *n* = 32 from *n* = 102 for restored plots) (Fig. 3 and 4), and were more abundant in the restored plots (degraded 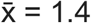; restored 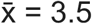; t = −2.9, df = 8, *p* = 0.01) (Fig. 3 and 4).

**Figure 3.**
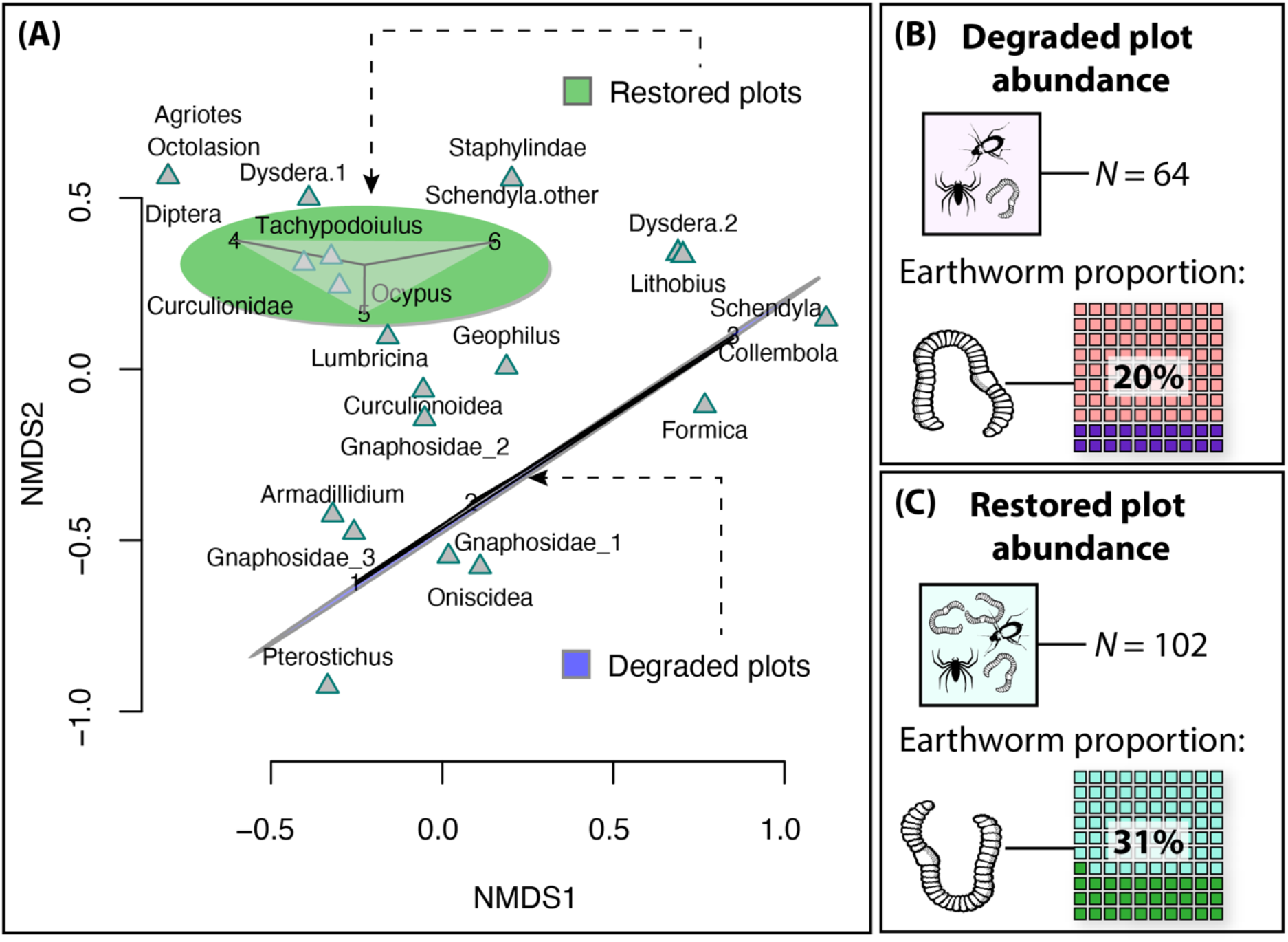
(A) Nonmetric multidimensional scaling (NMDS) ordination plots for visualising soil invertebrate beta diversity (community composition) for all plots (Stress: 0.01; Bray dissimilarity). Ellipses represent the standard error of the (weighted) average of scores. Clusters suggest clear differences between communities of the different treatment groups, as indicated by the colour purple ellipse for degraded plots (the linear ellipse) and green ellipse for restored plots. (B) Abundance of invertebrates counted in degraded plots and the proportion of earthworms. (C) Abundance of invertebrates counted in restored plots and the proportion of earthworms.

**Figure 4.**
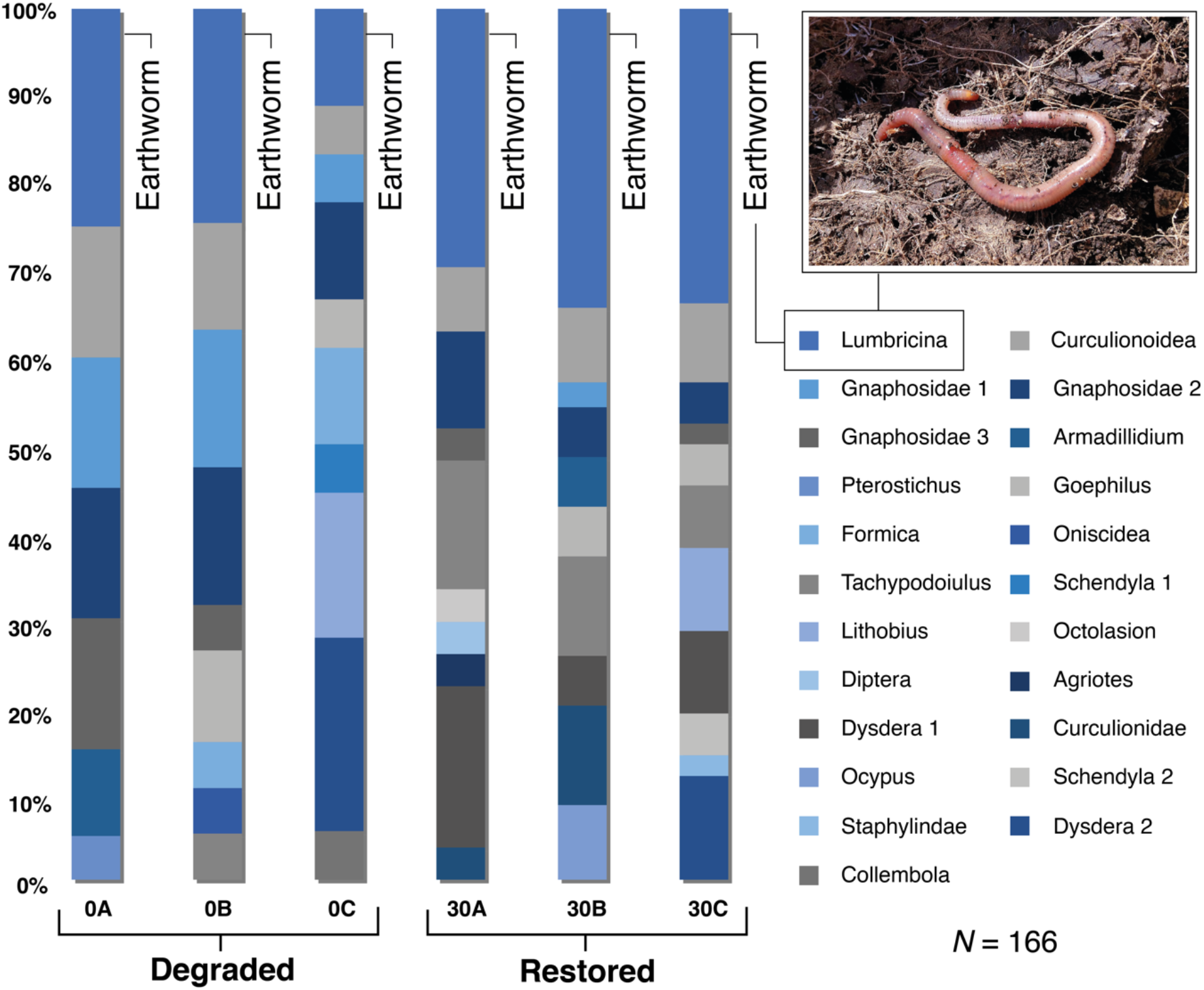
Stacked bar chart showing the relative abundance of soil invertebrates between plots (individual bars) and treatment groups (degraded vs restored). The top blue segment denotes earthworms (inset: earthworm), indicating a higher relative and absolute abundance of earthworms (*n* = 13 for degraded vs *n* = 32 for restored plots) in the samples from the restored forest plots.

#### Correlation of ecoacoustics variables and invertebrate abundance and richness

The ACI correlated with invertebrate abundance, with higher scores in the restored plots (Estimate = 0.2, R^2^ = 0.36, SE = 0.07, *p* = 0.01). A significant effect also occurred when changing ACI for BI (Estimate = 0.9, R^2^ = 0.31, SE = 0.03, *p* = 0.02). This suggests that restoration status and acoustic complexity and diversity metrics can predict invertebrate abundance. However, there was no significant effect of restoration/degradation or invertebrate richness on acoustic complexity based on the ACI (Estimate = −0.16, SE = 0.25, *p* = 0.5). This was also the case for acoustic diversity measured using the BI (Estimate = −1.63, SE = 1.18, *p* = 0.18). This corroborates the t-test for differences in means between invertebrate richness in the degraded vs restored plots (t = 0, df = 8, *p* = 1).

### Soil ecoacoustics in sound attenuation chamber

There was significantly greater ACI (Estimate = 1.6, R^2^ = 0.56, SE = 0.3, *p* = <0.01) (Fig. 5A) and BI (Estimate = 7.95, R^2^ = 0.58, SE = 1.8, *p* = <0.01) in restored compared with degraded soils, indicating bioacoustic complexity and diversity was higher in the restored plot soils in the sound attenuation chambers. However, there was no effect of restoration/degradation status on NDSI, indicating similar high-frequency to low-frequency ratios in the sound attenuation chamber for restored and degraded soils (Estimate = 0.04, SE = 0.13, *p* = 0.7) (t = −0.33, df = 8, *p* = 0.37) (Fig. 5A3).

**Figure 5.**
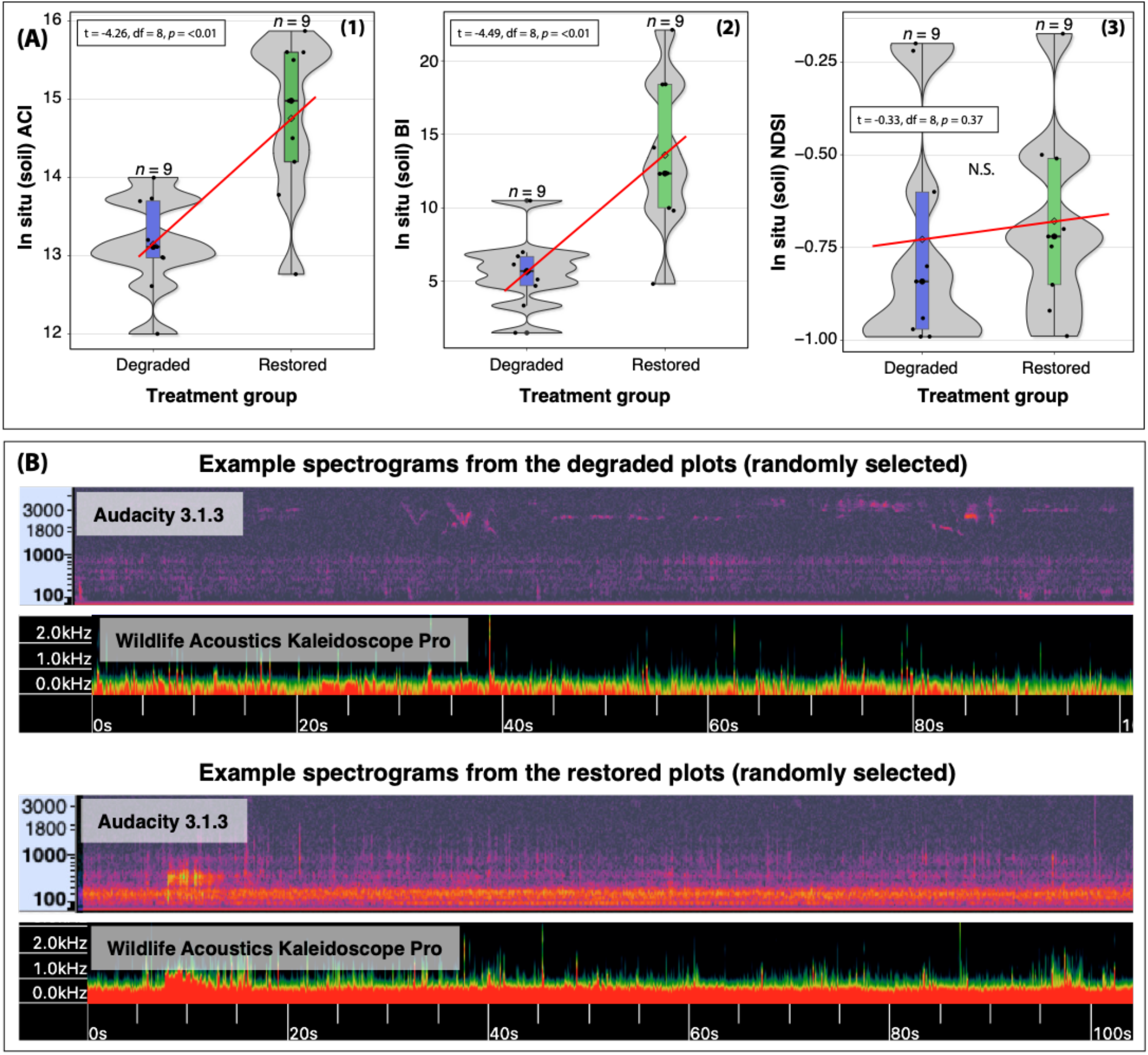
(A) Boxplots of acoustic index outputs for sound attenuation chamber (i.e., soil) samples and separated based on treatment groups (degraded vs restored). From left to right: (1) ACI, (2) BI, and (3) NDSI. Each plot has a red guideline to show trends in the mean values. (B) Examples of soil acoustic spectrogram for both treatment groups, showing the same window in two different analysis programmes (Wildlife Acoustics Kaleidoscope Pro and Audacity v3.1.3). N.S. = not significant.

### *In situ* soil ecoacoustics

There was no effect of the restoration/degradation status on ACI (Estimate = 0.12, SE = 0.14, *p* = 0.3) or BI (Estimate = 0.25, SE = 0.5, *p* = 0.6; Fig. 6). There was a greater NDSI in restored *in situ* soils than degraded soils (Estimate = 0.09, R^2^ = 0.15, SE = 0.03, *p* = 0.02) (t = −2.18, df = 89, *p* = 0.01) (Fig. 6, final plot), indicating greater high-frequency to low-frequency ratio in the restored soils.

**Figure 6.**
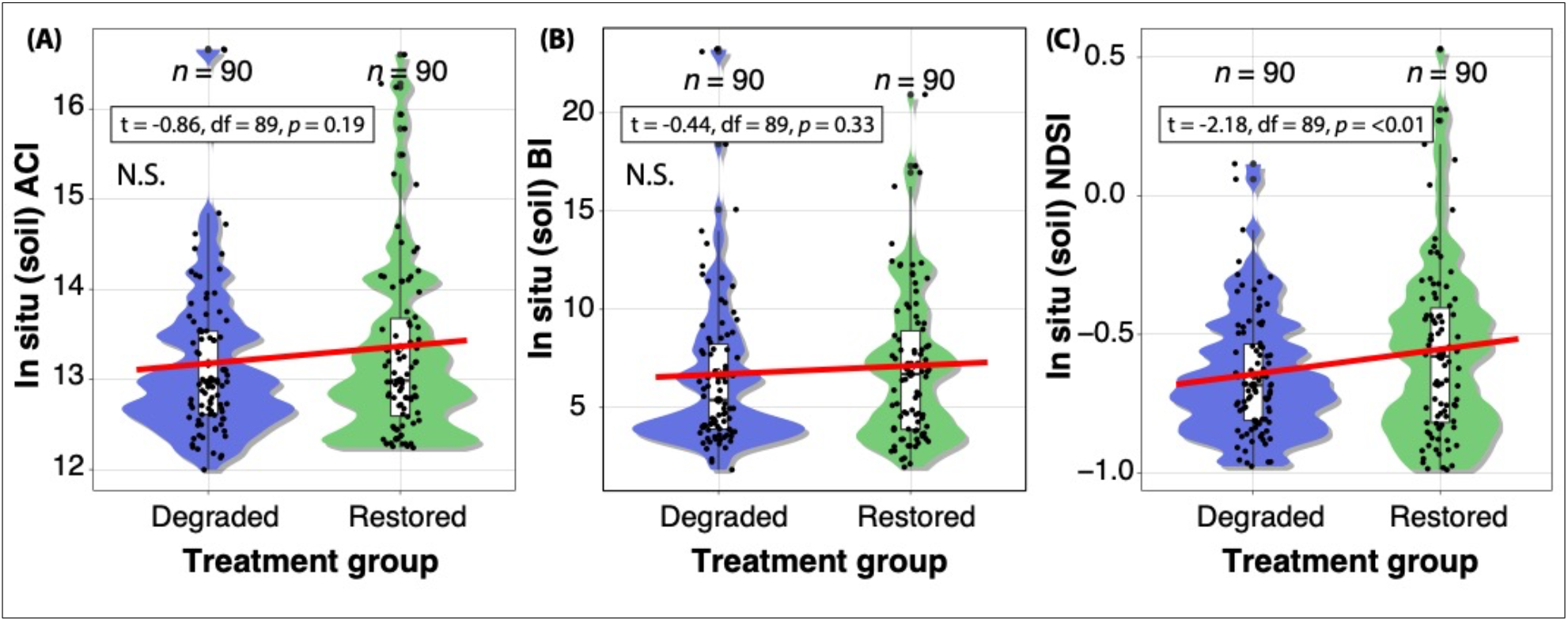
Boxplots of acoustic index outputs for *in situ* (i.e., soil) samples, separated by treatment group (degraded vs restored). From left to right: (A) ACI, (B) BI, and (C) NDSI. Each plot has a red guideline to show trends in the mean values. N.S. = not significant.

### Above-ground acoustic diversity and complexity

There was no effect of restoration/degradation status on ambient ACI (Estimate = − 0.5, SE = 0.6, *p* = 0.4) and BI (Estimate = 0.7, SE = 2.0, *p* = 0.7) (Fig. 7). When accounting for the visit and plot random effects, there was no effect of restoration/degradation status on ambient NDSI (Estimate = 0.14, SE = 0.2, *p* = 0.6).

**Figure 7.**
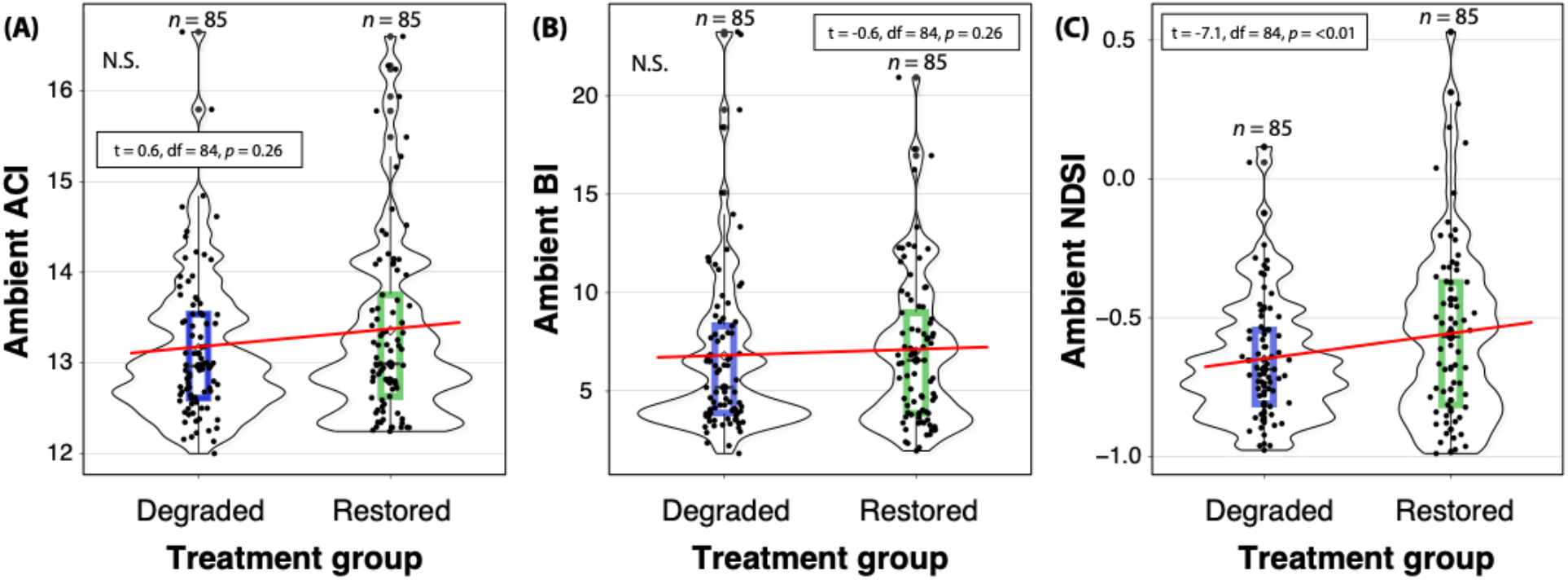
Boxplots of acoustic index outputs for ambient (i.e., above-ground) samples, separated by treatment group (degraded vs restored). From left to right: (A) ACI, (B) BI, and (C) NDSI. Each plot has a red guideline to show trends in the mean values. N.S. = not significant.

However, we do report a higher NDSI in the restored plots when we did a simple linear regression (Estimate = 0.18, R^2^ = 0.46, df = 168, *p* = 0.04).

## Discussion

We show that restored forest soils – in sound attenuation chambers at least – exhibit higher acoustic complexity and diversity than degraded soils, supporting our first hypothesis. Interestingly, there was no significant relationship between ambient (i.e., above-ground) acoustic diversity and degraded/restored status, probably in part due to the broad scale of sound transmission through the forest, compared to the highly localised soil soundscape (discussed below). We report greater high-frequency to low-frequency ratios in restored compared with degraded forest soils measured *in situ*, supporting our second hypothesis. Moreover, we validate our findings by reporting that invertebrate abundance – though not richness – was higher in restored than degraded forest soils. Accordingly, our study provides a case study on how soil ecoacoustics has clear potential to assess biodiversity in – and the restoration status of – forest soils.

### Restored vs. degraded soil ecoacoustics

Responses of soil biota to microhabitat conditions have been investigated extensively (Martins et al. 2012; Heiniger et al. 2015), and a recent study explored the temporal and spatial dynamics of soil biophony using ecoacoustics (Maeder et al. 2022). However, to our knowledge, our study is the first to investigate soil acoustic dynamics in a restoration context. It is the first study to relate the acoustic complexity, amplitude and frequency-band characteristics of the soil soundscape (via the ACI, BI and NDSI) to the abundance and richness of directly measured forest soil invertebrates. We reveal significant differences in the acoustic complexity and diversity between degraded and restored forest plots when measured in a sound attenuation chamber. These differences were associated with soil invertebrate abundance but not richness (unlike the findings of Maeder et al. 2022). This relationship between acoustic signals and soil communities, and the variation between degraded and restored plots, suggests that the restoration status of forest soils can be captured by monitoring soil soundscapes. Our models show that we could predict acoustic complexity and diversity based on the degraded and/or restored status of the forest plots, and these relationships were still significant when accounting for plot and visit-associated variability. The Acoustic Complexity Index (ACI) was the only one of the three indices we used that assesses the temporal dynamics of the sound recordings. It has become clear during this study that soil recordings are characterised by broadband stop-start intermittent noises produced by soil fauna, and these dynamics are better represented in the time domain than by analysing patterns across frequency bins (as done with BI and NDSI). Therefore, ACI is the best index to analyse this characteristic.

However, our results contrasted somewhat between samples from the sound attenuation chamber and taken *in situ*. The reason for this could be that the chamber may enhance the quality of the acoustic signal and reduce external noise. Despite the resting period, the act of moving soil into the chamber could also stimulate the movement (and hence sound production) of soil fauna, although acoustic complexity and diversity were still significantly higher in the restored soils. These findings suggest that the sound attenuation chamber sampling approach may be more suitable for detecting soil fauna acoustic signals in this forest restoration context. However, the *in situ* approach has the benefit of being less intrusive (i.e., no soil excavation is required). Therefore, it will be important to further optimise the *in situ* sampling strategy to improve the application of ecoacoustics to restoration.

The lack of association between soil invertebrate *richness* and acoustic index outputs contradicts the relationships found in a recent soil acoustics study (Maeder et al. 2022). This could simply be due to inter-ecosystem variability and the variety of acoustic signals made by soil fauna, which is still poorly understood. Alternatively, it could result from the relatively rapid *in situ* invertebrate-counting method employed in this study, which only provided a ‘snapshot’ of the resident soil fauna. Mean invertebrate richness was the same for both degraded and restored forest plots, although the invertebrate abundance was significantly higher in the restored plots. This aligns with other studies that show higher soil invertebrate abundance in habitats with lower disturbance (Smith et al. 2008; Nkem et al. 2020). The higher abundance of earthworms in the restored soils also corroborates other studies (Wodika et al. 2014; Singh et al. 2020). This could partially explain the higher acoustic complexity detected in restored soils. For instance, earthworms form burrows through the soil as they seek carbon-rich areas, which serve as preferential networking pathways for plant root growth, water flow and gas transport (Lacoste et al. 2018), all of which contribute to the soil soundscape (Gagliano et al. 2017; Del Stabile et al. 2022; Keen et al. 2022). In the future, it would be prudent to take a more robust approach to invertebrate counting, such as using the Berlese method (Sabu and Shiju 2010). This involves specially-adapted funnels to separate soil invertebrates from litter and particles and counting *ex situ* (Maeder et al. 2022). Metagenomics analysis is another option, either alone or in combination with traditional methods. This allows the genomes of soil organisms to be sequenced, differentiated and labelled without requiring morphological analysis (Schmidt et al. 2022). However, the need to control false-positive occurrences resulting from legacy DNA is vital (Laroche et al. 2017).

We report a significant association between NDSI values and the degradation/restoration status of forest plots, where restored plots exhibited a greater high-frequency to low-frequency ratio, aligning with our hypothesis. The NDSI seeks to describe the ‘health’ of an ecosystem by inferring the level of anthropogenic disturbance received (Eldridge et al. 2016). We hypothesised that our recording devices were more likely to detect higher-frequency biophony in restored plots and lower-frequency anthropogenic disturbance in degraded plots. This was based on the assumption that the increased signals from biological activity in restored plots would outweigh low-frequency noise, with potential effects also from the attenuation properties of the system (Tashakor and Chamani 2021; Sangermano 2022) i.e., the energy loss of sound propagation in a given medium. It could also be that greater earthworm activity changes soil characteristics (making them more air permeable) to allow better propagation of higher-frequency sounds, thereby increasing NDSI scores (Keen et al. 2022). Understanding the factors that affect this biophony-to-anthrophony ratio in a restoration context warrants further research. Examples of next steps could be conducting controlled experiments that manipulate sound sources and adding/removing vegetation and other physical features and media that provide noise attenuation. Applying new physics-based models to evaluate how the frequency and distance-dependent attenuation of sound impact the acoustic detection of soniferous species (Haupert et al. 2022) could also improve outcomes in a restoration monitoring context. Interestingly, there was no significant difference in the NDSI values between degraded and restored soil in the sound chambers, which was probably because the sound attenuation foam in the chamber acts to standardise ambient acoustic conditions.

### Above-ground ecoacoustics

Contrary to our expectations, we did not find a significant relationship between above-ground acoustic diversity and complexity and the degradation/restoration status of the forest plots. We hypothesised that we would observe higher acoustic diversity in the restored forest plots as faunal species richness, abundance, biomass and functional diversity are known to increase with restoration age (Derhé et al. 2016). Moreover, studies have shown that bird species diversity (the most soniferous group contributing to the soundscape) increases as restored forests mature, and bird communities in recovering areas become more similar to those of undisturbed areas with post-restoration age (Owen et al. 2021). The lack of a restoration effect on above-ground acoustic diversity and complexity could be due to our degraded and restored plots being relatively small compared to the soundscape of birdsong. Consequently, birdsong acoustic signals could potentially overlap across our plots, which is a limitation of our study. Future studies should pair sampling in time across plots, particularly when degraded and restored plots are within relatively close proximity to each other. Alternatively, mean acoustic diversity might increase as patch size increases, and more complex vegetation is associated with higher diversity (Grant et al. 2016). Therefore, it is possible that the minimum habitat patch size in our study was not sufficient to influence acoustic source variability in the treatment groups.

Our study provides preliminary evidence for using soil ecoacoustics – a minimally-intrusive and cost-effective assessment method – as a soil biota monitoring tool that can evaluate restoration projects. With future work, soil ecoacoustics could develop into an effective tool that measures the abundance, complexity and composition of soil biota that is also sensitive to restoration interventions. Given the rapid pace of biodiversity loss and the rise in anthropogenic noise, the ability to detect the acoustic signals from soniferous species and monitor the level of disturbance from anthrophonies has never been more important. Further exploration of above-ground ecoacoustics in different forest restoration settings, e.g., sites receiving different restoration interventions of varying patch sizes and in different biomes, would be valuable. Building on our findings—that soil acoustic complexity and diversity and noise disturbance differ between degraded and restored forest plots—has the potential to inform and enhance future restoration policy and practice.

## Literature Cited

Abrahams C (2019) Comparison between lek counts and bioacoustic recording for monitoring Western Capercaillie (Tetrao urogallus L.). Journal of Ornithology, 160: 685–697

Abrahams C, Geary M (2020) Combining bioacoustics and occupancy modelling for improved monitoring of rare breeding bird populations. Ecological Indicators, 112: 106131.

Abrahams C, Desjonquères C, Greenhalgh J (2021) Pond Acoustic Sampling Scheme: A draft protocol for rapid acoustic data collection in small waterbodies. Ecology and Evolution, 11:7532–7543

Adobe (2021) Adobe Illustrator. https://helpx.adobe.com/illustrator/using/whats-new.html (accessed 13 November 2022)

Bates D, Mächler M, Bolker B, Walker S (2015) Fitting Linear Mixed-Effects Models Using lme. Journal of Statistical Software. 67: 1–48.

Boelman NT, Asner GP, Hart PJ, Martin RE (2007) Multi-trophic invasion resistance in Hawaii: bioacoustics, field surveys, and airborne remote sensing. Ecological Applications 17:2137–2144.

Breed MF, Harrison PA, Blyth C, Byrne M, Gaget V, Gellie NJ, Groom SV, Hodgson R, Mills JG, Prowse TA, Steane DA (2019) The potential of genomics for restoring ecosystems and biodiversity. Nature Reviews Genetics, 20:615–628.

Camarretta N, Harrison PA, Bailey T, Potts B, Lucieer A, Davidson N, Hunt M (2020) Monitoring forest structure to guide adaptive management of forest restoration: a review of remote sensing approaches. New Forests, 51:573–596.

de Almeida DR, Stark SC, Valbuena R, Broadbent EN, Silva TS, de Resende AF, Ferreira MP, Cardil A, Silva CA, Amazonas N, Zambrano AM (2020) A new era in forest restoration monitoring. Restoration Ecology, 28:8–11.

de Framond L, Brumm H (2022) Long-term effects of noise pollution on the avian dawn chorus: a natural experiment facilitated by the closure of an international airport. Proceedings of the Royal Society B, 289:20220906.

De Jong K, Amorim MCP, Fonseca PJ, Fox CJ, Heubel KU (2018) Noise can affect acoustic communication and subsequent spawning success in fish. Environmental Pollution, 237:814–823.

De Vos JM, Joppa LN, Gittleman JL, Stephens PR, Pimm SL (2015) Estimating the normal background rate of species extinction. Conservation Biology, 29:452–462.

Del Stabile F, Marsili V, Forti L, Arru L (2022) Is There a Role for Sound in Plants?. Plants, 11:2391.

Derhé MA, Murphy H, Monteith G, Menéndez R (2016) Measuring the success of reforestation for restoring biodiversity and ecosystem functioning. Journal of Applied Ecology, 53:1714–1724.

Eldridge A, Casey M, Moscoso P Peck M (2016) A new method for ecoacoustics? Toward the extraction and evaluation of ecologically-meaningful soundscape components using sparse coding methods. PeerJ, 4:e2108.

Eldridge A, Guyot P, Moscoso P, Johnston A, Eyre-Walker Y, Peck M (2018) Sounding out ecoacoustic metrics: Avian species richness is predicted by acoustic indices in temperate but not tropical habitats. Ecological Indicators, 95:939–952.

Gamal MA, Khalil MH, Maher G (2020) Monitoring and Studying Audible Sounds Inside Different Types of Soil and Great Expectations for its Future Applications. Pure Applied Geophysics, 177:5397–5416.

Gan H, Zhang J, Towsey M, Truskinger A, Stark D, van Rensburg BJ, Li Y, Roe P (2020) Data selection in frog chorusing recognition with acoustic indices. Ecological Information, 60:101160.

Gagliano M, Grimonprez M, Depczynski M, Renton M (2017) Tuned in: plant roots use sound to locate water. Oecologia, 184:151–160.

Glatthorn J, Annighöfer P, Balkenhol N, Leuschner C, Polle A, Scheu S, Schuldt A, Schuldt B, Ammer C (2021) An interdisciplinary framework to describe and evaluate the functioning of forest ecosystems. Basic Applied Ecology, 52:1–14.

Golar G, Muis H, Akhbar A, Khaeruddin C (2022) Threat of Forest Degradation in Ex-Forest Concession Right (HPH) in Indonesia. Sustain Climate Change, 15:216–223.

Gollan JR, de Bruyn LL, Reid N, Wilkie L (2013) Monitoring the ecosystem service provided by dung beetles offers benefits over commonly used biodiversity metrics and a traditional trapping method. Journal of Nature Conservation, 21:183–188.

Grant BB, Samways MJ (2016) Use of ecoacoustics to determine biodiversity patterns across ecological gradients. Conservation Biology, 30:1320–1329.

Guidi C, Bou-Cabo M, Lara G, KM3NeT Collaboration (2021) Passive acoustic monitoring of cetaceans with KM3NeT acoustic receivers. Journal of Instruments, 16:C10004.

Hansen AJ, Noble BP, Veneros J, East A, Goetz SJ, Supples C, Watson JE, Jantz PA, Pillay R, Jetz W, Ferrier S (2021) Toward monitoring forest ecosystem integrity within the post-2020 Global Biodiversity Framework. Conservation Letters, 14:e12822.

Harvey DJ, Hawes CJ, Gange AC, Finch P, Chesmore D, Farr IAN (2011) Development of non-invasive monitoring methods for larvae and adults of the stag beetle, Lucanus cervus. Insect Conservation Diversity, 4:4–14.

Hastie GD, Lepper P, McKnight JC, Milne R, Russell DJ, Thompson D (2021) Acoustic risk balancing by marine mammals: anthropogenic noise can influence the foraging decisions by seals. Journal of Applied Ecology, 58:1854–1863.

Haupert S, Sèbe F, Sueur J (2022) Physics-based model to predict the acoustic detection distance of terrestrial autonomous recording units over the diel cycle and across seasons: Insights from an Alpine and a Neotropical forest. Methods in Ecology Evolution.

Heiniger C, Barot S, Ponge JF, Salmon S, Meriguet J, Carmignac D, Suillerot M Dubs F (2015) Collembolan preferences for soil and microclimate in forest and pasture communities. Soil Biology and Biochemistry, 86:181–192.

Hintze F, Machado RB, Bernard E (2021) Bioacoustics for in situ validation of species distribution modelling: An example with bats in Brazil. PLOS ONE, 16:e0248797.

Jones B, Zapetis M, Samuelson MM, Ridgway S (2020) Sounds produced by bottlenose dolphins (Tursiops): A review of the defining characteristics and acoustic criteria of the dolphin vocal repertoire. Bioacoustics, 29:399–440.

Kassambara A. (2022) Rstatix package. https://cran.r-project.org/web/packages/rstatix/index.html (accessed 10th November 2022).

Kasten P, Stuart HG, Jordan F, and Wooyeong J (2012) The Remote Environmental Assessment Laboratory’s Acoustic Library: An Archive for Studying Soundscape Ecology. Ecological Information 12:50–67.

Keen SC, Wackett AA, Willenbring JK, Yoo K, Jonsson H, Clow T, Klaminder J (2022) Non-native species change the tune of tundra soils: Novel access to soundscapes of the Arctic earthworm invasion. Science of the Total Environment, 838:155976.

Kleist NJ, Guralnick RP, Cruz A, Lowry CA Francis CD (2018) Chronic anthropogenic noise disrupts glucocorticoid signaling and has multiple effects on fitness in an avian community. Proceedings of the National Academy of Sciences, 115:E648–E657.

Kimmins JP (2004) Forest ecology. Fishes and forestry: Worldwide watershed interactions and management, 17–43.

Kissling WD, Ahumada JA, Bowser A, Fernandez M, Fernández N, García EA, Guralnick RP, Isaac NJ, Kelling S, Los W, McRae L (2018) Building essential biodiversity variables (EBVs) of species distribution and abundance at a global scale. Biological Reviews, 93:600–625.

Kühn S, Utne-Palm AC de Jong K (2022) Two of the most common crustacean zooplankton Meganyctiphanes norvegica and Calanus spp. produce sounds within the hearing range of their fish predators. Bioacoustics, 1–17.

Kuznetsova A (2020) The LmerTest package in R. https://cran.r-project.org/web/packages/lmerTest/index.html (accessed on 10th November 2022).

Lacoste M, Ruiz S Or D (2018) Listening to earthworms burrowing and roots growing-acoustic signatures of soil biological activity. Science Reports, 8:1–9.

Laroche O, Wood SA, Tremblay LA, Lear G, Ellis JI Pochon X (2017) Metabarcoding monitoring analysis: the pros and cons of using co-extracted environmental DNA and RNA data to assess offshore oil production impacts on benthic communities. PeerJ, 5:e3347.

Le Bayon RC, Bullinger G, Schomburg A, Turberg P, Brunner P, Schlaepfer R, Guenat C (2021) Earthworms, plants, and soils. Hydrogeol, Chem Weather, Soil Form, 81–103.

Li RQ, Dong M, Cui JY, Zhang LL, Cui QG, He WM (2007) Quantification of the impact of land-use changes on ecosystem services: a case study in Pingbian County, China. Environmental Monitoring Assessment, 128:03–510.

López-Baucells A, Yoh N, Rocha R, Bobrowiec PE, Palmeirim JM, Meyer CF (2021) Optimizing bat bioacoustic surveys in human-modified Neotropical landscapes. Ecological Applications, 31:e02366.

Maeder M, Gossner MM, Keller A, Neukom M (2019) Sounding soil: An acoustic, ecological artistic investigation of soil life. Soundscape J, 18:005–014.

Maeder M, Guo X, Neff F, Schneider Mathis D, Gossner MM (2022). Temporal and spatial dynamics in soil acoustics and their relation to soil animal diversity. PLOS ONE, 17:e0263618.

Mankin R (2022) Subterranean Arthropod Biotremology: Ecological and Economic Contexts. In P. S. M. Hill, V. Mazzoni, N. Stritih-Peljhan, M. Virant-Doberlet, & A. Wessel (Eds.), Biotremology: Physiology, Ecology, and Evolution, 8:511–527. Springer International Publishing, New York.

Martins da Silva P, Berg MP, Serrano AR, Dubs F, Sousa JP (2012) Environmental factors at different spatial scales governing soil fauna community patterns in fragmented forests. Landscape Ecology, 27:1337–1349.

Messier C, Bauhus J, Doyon F, Maure F, Sousa-Silva R, Nolet P, Mina M, Aquilué N, Fortin MJ, Puettmann K (2019) The functional complex network approach to foster forest resilience to global changes. Forest Ecosystems, 6:1–16.

Müller FG (2000) Does the convention on biodiversity safeguard biological diversity?. Environmental Values, 9:55–80.

Müller S, Gossner MM, Penone C, Jung K, Renner SC, Farina A, Anhäuser L, Ayasse M, Boch S, Haensel F, Heitzmann J (2022) Land-use intensity and landscape structure drive the acoustic composition of grasslands. Agriculture, Ecosystems Environment, 328:107845.

Nkem JN, Lobry de Bruyn L, King K (2020) The Effect of Increasing Topsoil Disturbance on Surface-Active Invertebrate Composition and Abundance under Grazing and Cropping Regimes on Vertisols in North-West New South Wales, Australia. Insects, 11:237.

O’Connor B, Bojinski S, Röösli C, Schaepman ME (2020) Monitoring global changes in biodiversity and climate essential as ecological crisis intensifies. Ecological Informatics, 55:101033.

Oksanen J, Simpson GL, Blanchet G, Kindt R, Legendre P, Minchin PR, O’Hara RB, Solymos P, Stevens H, Szoecs E, Wagner H, Barbour M, Bedward M, Bolker B, Borcard D, Carvalho G, Chirico M, Caceres MD, Duran S, Evangelista HBA, FitzJohn R, Friendly M, Furneaux B, Hannigan G, Hill MO, Lahti L, McGlinn D, Ouellette MH, Cunha ER, Smith T, Stier A, Braak CJFT, Weedon J (2022) The Vegan community ecology package in R. https://cran.r-project.org/web/packages/lmerTest/index.html (accessed on 10th November 2022).

Owen K, Mennill DJ, Campos FA, Fedigan LM, Gillespie TW, Melin AD (2020) Bioacoustic analyses reveal that bird communities recover with forest succession in tropical dry forests. COPA. http://dx.doi.org/10.5751/ACE-01615-150125

Pardos M, Del Río M, Pretzsch H, Jactel H, Bielak K, Bravo F, Brazaitis G, Defossez E, Engel M, Godvod K, Jacobs K (2021) The greater resilience of mixed forests to drought mainly depends on their composition: Analysis along a climate gradient across Europe. Forest Ecology Management, 481:118687.

Pieretti N, Farina A, Morri D (2011) A new methodology to infer the singing activity of an avian community: The Acoustic Complexity Index (ACI). Ecological Indicators, 11:868–873.

Popper AN, Hawkins AD (2019) An overview of fish bioacoustics and the impacts of anthropogenic sounds on fishes. Journal of Fish Biology, 94:692–713.

R Core Team (2022) R: A language and environment for statistical computing. R Foundation for Statistical Computing, Vienna, Austria. https://www.R-project.org/ (accessed on 10^th^ November 2022).

Robinson JM, Cameron R, Parker B (2021) The effects of anthropogenic sound and artificial light exposure on microbiomes: ecological and public health implications. Frontiers in Ecology and Evolution, 9:662588.

Robinson JM, Watkins H, Man I, Liddicoat C, Cameron R, Parker B, Cruz M, Meagher L (2021) Microbiome-Inspired Green Infrastructure: a bioscience roadmap for urban ecosystem health. ARQ. 25:292–303.

Robinson JM, Harrison PA, Mavoa S, Breed MF (2022) Existing and emerging uses of drones in restoration ecology. Methods in Ecology Evolution, 13:1899–1911.

Rodwell JS, Joint nature conservation committee (GB) (2006) National vegetation classification: Users’ handbook. Joint nature conservation committee, Peterborough.

Sabu TK, Shiju RT (2010) Efficacy of pitfall trapping, Winkler and Berlese extraction methods for measuring ground-dwelling arthropods in moist deciduous forests in the Western Ghats. Journal of Insect Science, 10.

Sama S (2021) Strengthening the Role of Forests in Climate Change Mitigation through the European Union Forest Law Enforcement, Governance and Trade Action Plan. Journal of Environmental Law and Policy, 1:1.

Sangermano F (2022) Acoustic diversity of forested landscapes: Relationships to habitat structure and anthropogenic pressure. Landscape and Urban Planning, 226:104508.

Schmidt A, Schneider C, Decker P, Hohberg K, Römbke J, Lehmitz R, Bálint M (2022) Shotgun metagenomics of soil invertebrate communities reflects taxonomy, biomass, and reference genome properties. Ecology and Evolution, 12:e8991.

Seidl R, Turner MG (2022) Post-disturbance reorganization of forest ecosystems in a changing world. Proceedings of the National Academy of Sciences, 119:e2202190119.

Simard SW (2018) Mycorrhizal networks facilitate tree communication, learning, and memory. In Memory and learning in plants, 191–213. Springer, Cham.

Singh S, Sharma A, Khajuria K, Singh J, Vig AP (2020). Soil properties changes earthworm diversity indices in different agro-ecosystem. BMC Ecology, 20:1–14.

Smith RG, McSwiney CP, Grandy AS, Suwanwaree P, Snider RM, Robertson GP (2008) Diversity and abundance of earthworms across an agricultural land-use intensity gradient. Soil Tilling Research, 100:83–88.

Stanturf JA, Palik BJ, Williams MI, Dumroese RK, Madsen P (2014) Forest restoration paradigms. Journal of Sustainable Forestry, 33:S161–S194.

Stoddard MT, McGlone CM, Fulé PZ, Laughlin DC, Daniels ML (2011) Native plants dominate understory vegetation following ponderosa pine forest restoration treatments. Western North American Naturalist, 71:206–214.

Stowell D, Sueur J (2020) Ecoacoustics: acoustic sensing for biodiversity monitoring at scale. Remote Sensing in Ecology and Conservation, 6:217–219.

Stroud JL (2019) Soil health pilot study in England: Outcomes from an on-farm earthworm survey. PLOS ONE, 14:e0203909.

Sueur J, Aubin T, Simonis C (2008) Seewave, a free modular tool for sound analysis and synthesis. Bioacoustics, 18:213–226.

Tashakor S, Chamani A (2021) Temporal variability of noise pollution attenuation by vegetation in urban parks. Environmental Science and Pollution Research, 28:23143–23151.

Taye FA, Folkersen MV, Fleming CM, Buckwell A, Mackey B, Diwakar KC, Le D, Hasan S, Saint Ange C (2021) The economic values of global forest ecosystem services: A meta-analysis. Ecological Economics, 189:107145.

Teixeira D, Maron M, van Rensburg BJ (2019) Bioacoustic monitoring of animal vocal behavior for conservation. Conservation Science and Practice, 1:e72.

Tidau S, Briffa M (2019) Anthropogenic noise pollution reverses grouping behaviour in hermit crabs. Animal Behavior, 151:113–120.

Turner A, Fischer M, Tzanopoulos J (2018) Sound-mapping a coniferous forest— Perspectives for biodiversity monitoring and noise mitigation. PLOS ONE, 13:e0189843.

Yu Y, Zhao W, Martinez-Murillo JF, Pereira P (2020) Loess Plateau: from degradation to restoration. Science of the Total Environment, 738:140206.

van der Mescht AC, Pryke JS, Gaigher R, Samways MJ (2021) Ecological and acoustic responses of bush crickets to anthropogenic and natural ecotones. Biodiversity Conservation, 30:3859–3878.

Vega-Hidalgo Á, Flatt E, Whitworth A, Symes L (2021) Acoustic assessment of experimental reforestation in a Costa Rican rainforest. Ecological Indicators, 133:108413.

Watson JE, Evans T, Venter O, Williams B, Tulloch A, Stewart C, Thompson I, Ray JC, Murray K, Salazar A, McAlpine C (2018) The exceptional value of intact forest ecosystems. Nature Ecology and Evolution, 2:599–610.

Wildlife Acoustics (2022) Kaleidoscope Pro sound analysis software. https://www.wildlifeacoustics.com/products/kaleidoscope-pro (accessed on 10th November 2022).

Williams-Linera G, Bonilla-Moheno M, López-Barrera F, Tolome J (2021) Litterfall, vegetation structure and tree composition as indicators of functional recovery in passive and active tropical cloud forest restoration. Forest Ecology and Management, 493:119260.

Wodika BR, Klopf RP, Baer SG (2014) Colonization and recovery of invertebrate ecosystem engineers during prairie restoration. Restoration Ecology, 22:456–464.

Zipkin EF, Zylstra ER, Wright AD, Saunders SP, Finley AO, Dietze MC, Itter MS Tingley MW (2021) Addressing data integration challenges to link ecological processes across scales. Frontiers in Ecology and the Environment, 19:30–38.

