## Supplementary materials for "The sound of restored soil: Measuring soil biodiversity in a forest restoration chronosequence with ecoacoustics"

### S1 Sampling duration pilot study

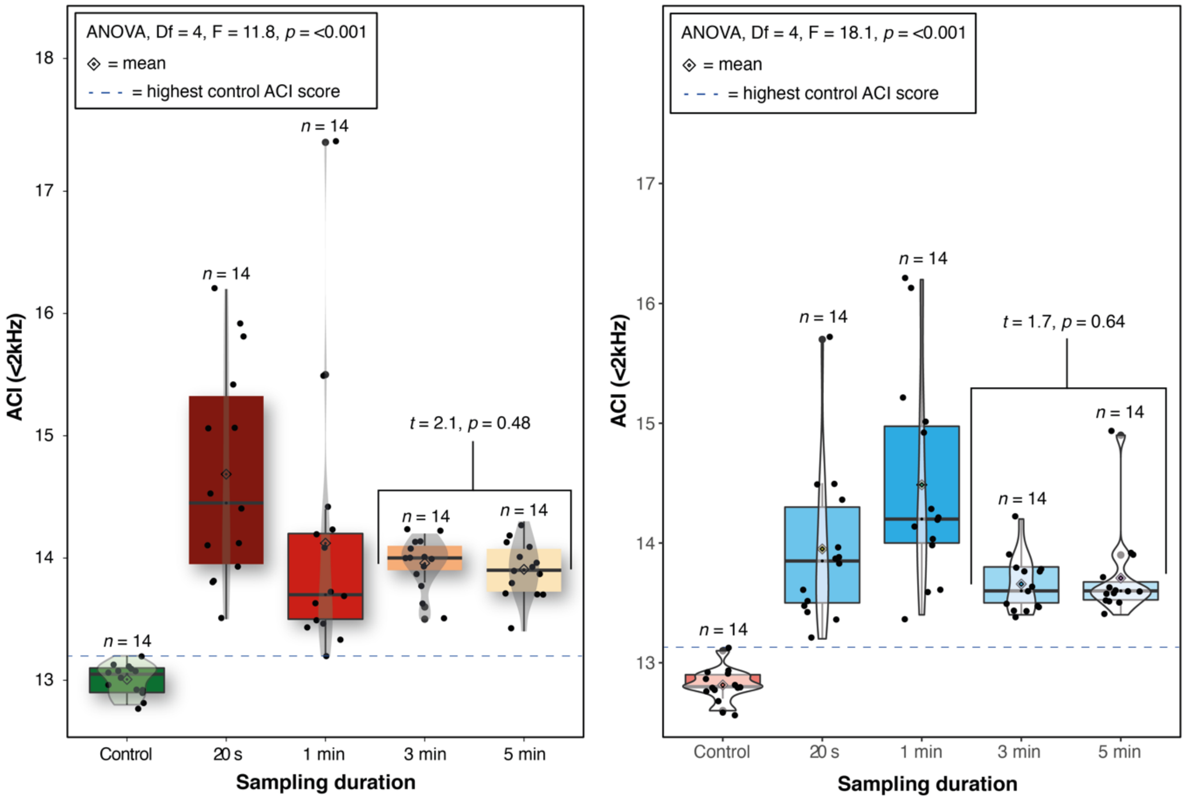


**Fig. S1.** Results of the Acoustic Complexity Index saturation test. Variability was high during the 20 s and 1 min sampling durations and plateaued at 3 mins.

### S2 Sound attenuation chamber tests

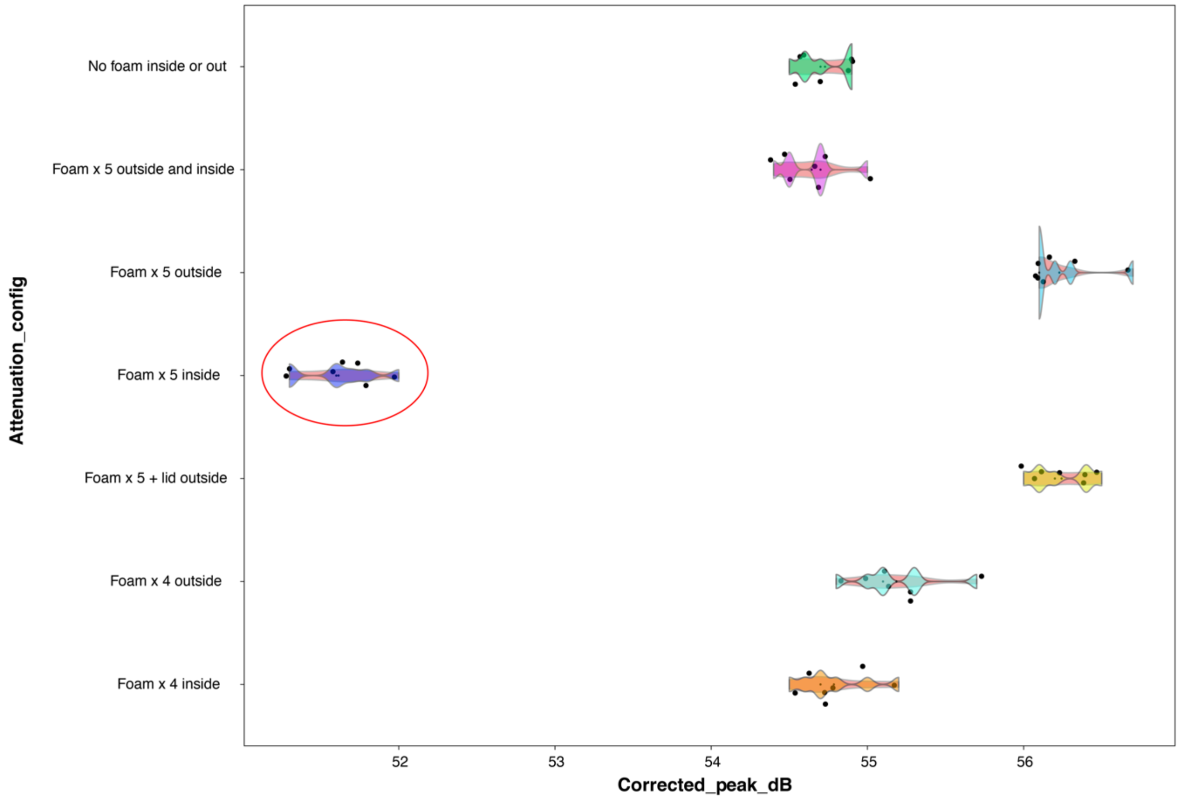


**Figure S2.** Boxplots showing sound attenuation tests, indicating the optimal performance of five foam inserts (in red circle).

**Table S1.** Sampling dates and weather conditions.

| Visit (date) | Ambient conditions |
|---|---|
| 1 (21st-23rd June 2022) | x̄ temperature: 23°C; Dry; Wind: Beaufort scale 1; x̄ relative humidity: 47.8% |
| 2 (26th-27th July 2022) | x̄ temperature: 21°C; Dry; Wind: Beaufort scale 1; x̄ relative humidity: 67.3% |
| 3 (11th August 2022) | x̄ temperature: 26°C, Dry; Wind: Beaufort scale 1; x̄ relative humidity: 54.6% |
